# A modified porous silicon microparticle promotes mucosal delivery of SARS-CoV-2 antigen and induction of potent and durable systemic and mucosal T helper 1 skewed protective immunity

**DOI:** 10.1101/2021.11.22.469576

**Authors:** Awadalkareem Adam, Qing Shi, Binbin Wang, Jing Zou, Junhua Mai, Samantha R Osman, Wenzhe Wu, Xuping Xie, Patricia V Aguilar, Xiaoyong Bao, Pei-Yong Shi, Haifa Shen, Tian Wang

## Abstract

Development of optimal SARS-CoV-2 vaccines to induce potent, long-lasting immunity and provide cross-reactive protection against emerging variants remains a high priority. Here, we report that a modified porous silicon microparticle (mPSM)-adjuvanted SARS-CoV-2 receptor-binding domain (RBD) vaccine activated dendritic cells and generated more potent and durable SARS-CoV-2-specific systemic humoral and type 1 helper T (Th) cell-mediated immune responses than alum-formulated RBD following parenteral vaccination, and protected mice from SARS-CoV-2 and Beta variant infection. mPSM facilitated the uptake of SARS-CoV-2 RBD antigens by nasal and airway epithelial cells. Parenteral and intranasal prime and boost vaccinations with mPSM-RBD elicited potent systemic and lung resident memory T and B cells and SARS-CoV-2 specific IgA responses, and markedly diminished viral loads and inflammation in the lung following SARS-CoV-2 Delta variant infection. Our results suggest that mPSM can serve as potent adjuvant for SARS-CoV-2 subunit vaccine which is effective for systemic and mucosal vaccination.

## INTRODUCTION

The coronavirus disease 2019 (COVID-19) pandemic, which was caused by severe acute respiratory syndrome coronavirus 2 (SARS-CoV-2), has caused a devastating impact on global public health and economy over the past two years. SARS-CoV-2 belongs to the genus *Betacoronavirus* (β-COV) of the family *Coronaviridae* and contains a single-stranded positive-sense RNA genome. The genome encodes structural proteins (spike [S], envelope [E], membrane [M] and nucleocapsid [N]), nonstructural proteins (nsp1-nsp16), and several accessory proteins ^1^. The S protein is the major virus surface glycoprotein that engages the interaction with human angiotensin-converting enzyme 2 (hACE2) through its receptor-binding domain (RBD) and facilitates virus entry into target cells. Both the S protein and the RBD can elicit highly potent neutralizing antibodies (NAbs) and contain major T cell epitopes, thus have been the main targets for vaccine development ^2-4^.

In response to the pandemic, many vaccine platforms have been rapidly developed and tested to enable production of effective vaccines against SARS-CoV-2 infection. This includes inactivated vaccines, subunit vaccines, DNA vaccines, mRNA vaccines, viral vectored vaccines, and live-attenuated vaccines ^1,5-9^. Currently, three vaccines have been granted emergency use authorization (EUA) from the FDA. However, the increasing rate of emergence of variants with enhanced viral transmission and disease severity in COVID-19 patients ^10,11^, potential concerns of “vaccine-induced disease enhancement” ^12^ and risk of antibody-dependent enhancement due to waning immunity after vaccination ^13^ have together posed additional challenges for the global vaccine efficiency efforts. It is clear that continuous efforts toward optimizing existing vaccine platforms and development of more effective novel vaccines are needed.

In this study, we tested the immunogenicity of a novel adjuvant comprised of a modified porous silicon microparticle (mPSM) for the SARS-CoV-2 S protein RBD subunit vaccine (mPSM-RBD). We also assessed the protective efficacy of mPSM-RBD vaccine in animal models of SARS-CoV-2 infection. PSMs can serve as a carrier and a reservoir to maintain sustained release of proteins and peptide antigens inside dendritic cell (DC)s ^14^. We previously identified PSM as a potent activator of type I interferon (IFN I) responses in DCs, and its protective effects as an adjuvant for cancer vaccines to stimulate T helper 1 (Th1) immunity. More recently, we found that mPSM, prepared by loading the TLR9 ligand cytosine guanosine dinucleotide (CpG) and STING agonist cyclic 2’,3’-GAMP (cGAMP)-to PSMs, can serve as a more potent adjuvant for tumor antigen to elicit higher levels of IFN I and inflammatory cytokines in DCs than PSM alone, and induces strong anti-tumor Th1 type immunity^15^. Here, we report that mPSM-RBD vaccine triggers more potent, and durable systematic Th1-prone immune responses than alum-RBD following parenteral vaccination in mice and protects mice against SARS-CoV-2 and Beta variant infection. In addition, mPSM facilitated mucosal delivery of SARS-CoV-2 RBD antigens. Parenteral and mucosal prime-boost vaccination promoted the induction of SARS-CoV-2-specific systematic and lung-resident Th1 and IgA immune responses, and protected mice from SARS-CoV-2 Delta variant infection.

## RESULTS

### mPSM is a potent adjuvant for SARS-CoV-2 RBD subunit vaccine and triggers SARS-CoV-2 - specific antibody production with minimal adverse effects upon parenteral vaccination in mice

The RBD of SARS-CoV-2 S protein is considered to be the major protective antigen, which elicits highly potent neutralization antibodies ^4^. To express and purify the S RBD domain, a DNA fragment encoding amino acid residues 319 to 541 of SARS-CoV-2 S protein was cloned into the lentivirus vector pCDH-CMV-MCS-EF1α-RFP which was then applied to transduce 293FT cells. To facilitate the secretion and purification of the protein, the first 19 residues of the S protein and a hexahistidine (6xHis) tag were fused at the N-terminal as a secretion signal and the C-terminal respectively. The recombinant RBD protein (25 to 30 kDa) was purified from the cell culture supernatant (**Fig 1A-B**). The protein antigen was packaged into mPSM to prepare a SARS-CoV-2 RBD subunit vaccine (mPSM-RBD) following our recently described protocol ^15^. To assess the effects of mPSM-RBD on DC activation and antigen presentation, bone marrow-derived DCs (BMDCs) isolated from BALB/c mice were treated with PBS (mock), RBD alone or together with either Alum, or mPSM. The production of proinflammatory cytokines, including IL-6, IL-12p70, and TNF-α was markedly increased in mPSM-RBD -treated but not in alum-RBD-or mock -treated DCs. Cell surface co-stimulation molecules, such as CD80 and CD86 expression was also enhanced in the mPSM-RBD -treated, but not in the alum-RBD- treated DCs (**Fig 1C-D, Fig. S1A**), which together suggest a role of mPSM in promoting activation of antigen presenting cells (APC). To assess whether mPSM-RBD vaccination produces SARS-CoV-2-specific antibody responses, sera of mice vaccinated with RBD alone, alum-RBD, or mPSM-RBD were collected one month post vaccination to determine their inhibitory effects on RBD binding to its receptor ACE2. While serum from Alum-RBD-vaccinated mice diminished RBD binding to ACE2, that from mPSM-RBD-treated mice nearly abolished binding of ACE2 to RBD protein (**Fig 1E**). Routes of parenteral vaccination were also compared. Mice were primed and boosted with mPSM-RBD (5 μg) via intradermal (i.d.), intramuscular (i.m.), or intraperitoneal (i.p.) inoculation. All three routes of inoculation resulted in high titers of RBD-binding IgG2a, IgG2b, and IgG1 subtypes IgG antibodies at one month post vaccination (**Fig 1F**). To further assess the effects of mPSM-RBD dosing in mice, mice were vaccinated i.d. with 1 to 50 μg mPSM-RBD. Interestingly, vaccination with as little as 5 μg mPSM-RBD triggered similar levels of IgG2b responses as elicited by 25 and 50 μg mPSM-RBD, which remained high more than 180 days post vaccination. However, 25 and 50 μg mPSM-RBD triggered much stronger IgG2a and IgG1 responses than the 5 μg mPSM-RBD group (**Fig 1G**). Lastly, mPSM-RBD was applied to evaluate potential toxicity, and biomarkers including urea nitrogen (BUN), albumin (ALB), calcium (CA), creatinine (CRE), glucose (GLU), phosphorus (PHOS), and total protein (TP) were assessed. No significant difference between mPSM-RBD and PBS control was observed (**Fig S1B-E**), which indicates no severe toxicity from mPSM-RBD in mice. Overall, these results suggest that mPSM serves as a potent and safe adjuvant for SARS-CoV-2 RBD subunit vaccine.

**Figure 1.**
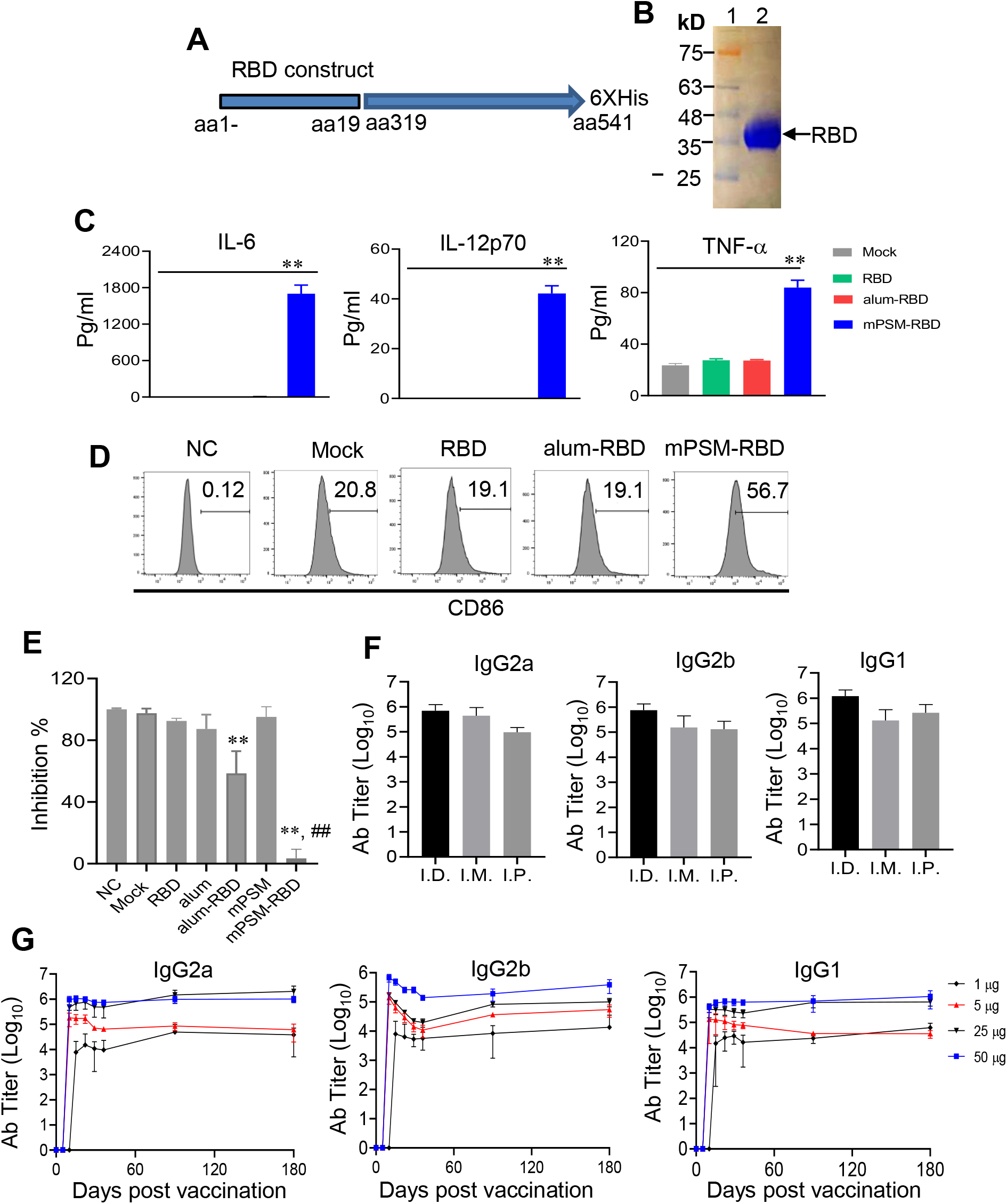
mPSM serves a potent adjuvant for SARS-CoV-2 RBD vaccine to generate SARS-CoV-2 specific antibodies in mice following parenteral vaccination. **A.** Schematic of SARS-CoV-2 RBD construct. **B.** Coomassie blue staining of purified recombinant (r)RBD protein. Lane 1: protein molecular weight marker. **C-D.** Cytokine production and activation of cell surface CD86 expression in BMDCs treated with mPSM-RBD and controls. **C.** Levels of IL-6, IL-12p70 and TNF-*α* in cell culture supernatant were determined by ELISA 24 h after the treatment. n = 3. **D.** CD86 expression was measured by flow cytometry analysis. One representative image was shown. **E.** ACE2 competition assay. Sera of mice-vaccinated with mPSM-RBD, alum-RBD, RBD, and mock were collected at 1 month post vaccination to measure the inhibitory effects on RBD binding to its receptor ACE2. n= 3-4. **F.** Endpoint IgG subtypes titers against SARS-CoV-2 RBD measured in sera collected 1 month post parenteral prime (day 0) and boost (day 14) vaccination. n =4. **G.** Endpoint IgG subtype titers against SARS-CoV-2 RBD measured in sera of mice following prime (day 0) and boost (day 14) vaccination with different doses of r-RBD-formulated with mPSM. n =3. ** *P* < 0.01 compared to mock group. ##*P* < 0.01 compared to alum-RBD group.

### Parenteral vaccination with mPSM-RBD subunit vaccine generated strong and durable systemic SARS-CoV-2-specific humoral and type 1 helper T (Th) cell-mediated immune responses in different strains of mice

BALB/c and C57BL/6 mice were i.d. inoculated with PBS control, RBD (5 μg) alone, alum-RBD (5 μg), or mPSM-RBD (5 μg) on day 0 and boosted with the same dose on day 14. Sera were collected at days 7, 14 and 21 to determine antibody titers (**Fig 2A**). mPSM-RBD group showed 10^3^ to 10^7^ titers of RBD binding IgG subtype antibodies (IgG2a, IgG2b, and IgG1) on days 7, 14 and 21. In comparison, alum-RBD vaccination barely induced any RBD IgG2a and IgG2b antibodies, and only low titers of RBD-binding IgG1 antibodies after day 14 (**Fig 2B**). While both alum-RBD and mPSM-RBD produced similar levels of RBD -binding IgG1 antibodies in B6 mice, only the latter induced RBD-specific IgG2b responses (**Fig S2A-B**). On day 30, mPSM-RBD -vaccinated BALB/c mice had over 3 -fold more SARS-COV-2 S-specific IgG^+^ splenic B cells (**Fig 2C, D**) and the splenocytes produced over 8 -fold higher IFN-γ upon *in vitro* re-stimulation with S peptide pools compared to the alum-RBD group (**Fig 2E, F**). mPSM-RBD vaccination also triggered more robust SARS-COV-2-specific splenic B and T cell responses in B6 mice compared to alum-RBD vaccine (**Fig S2C-F**). Cytokines secreted by Th1 cells are known to mediate isotype switching to IgG2a, whereas cytokines secreted by Th2 cells mediate isotype switching to IgG_1_ ^16^. Thus, the above results suggest that the mPSM-RBD vaccine promotes stronger humoral and Th1-prone immune responses in mice.

**Figure 2.**
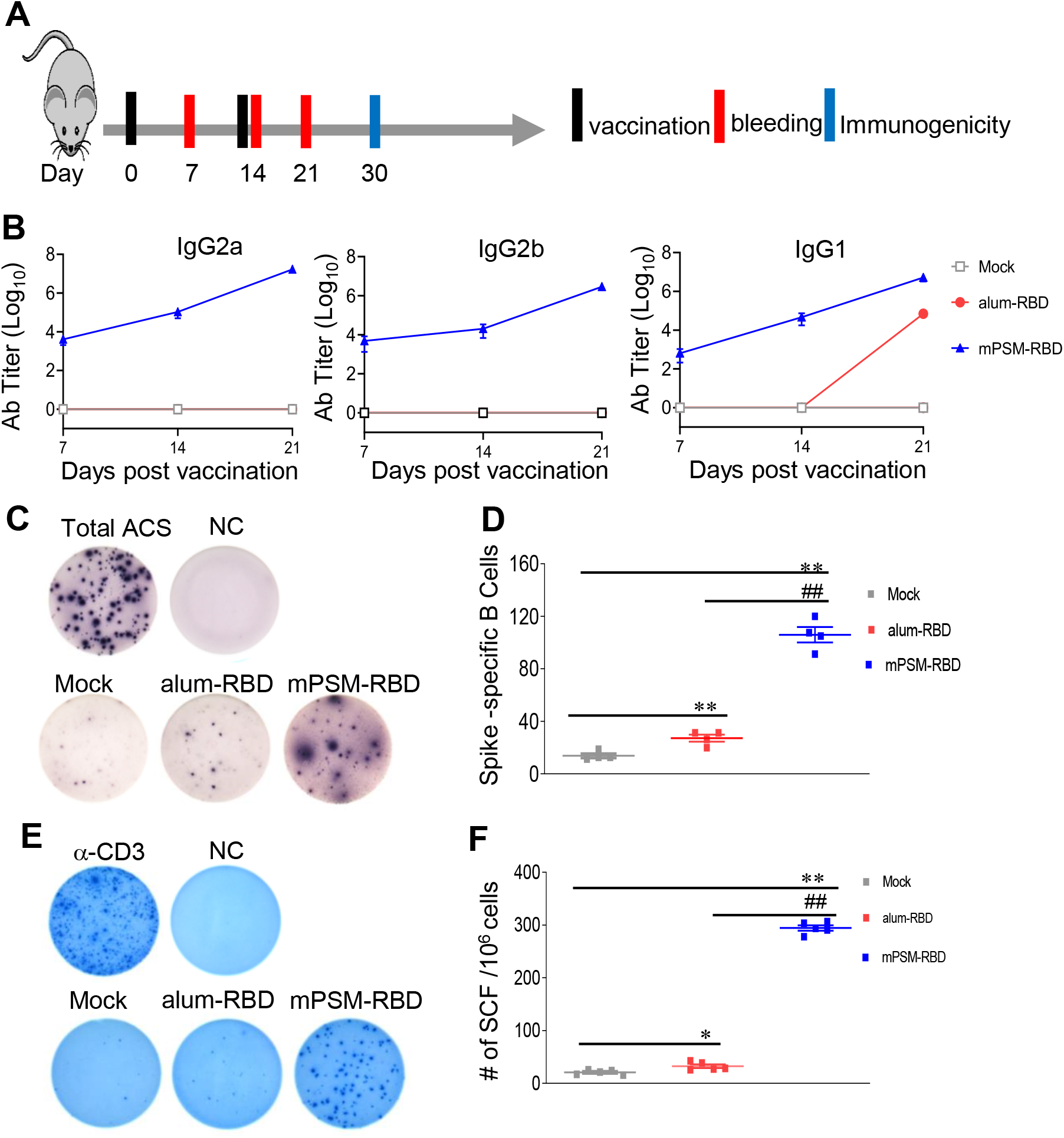
mPSM-RBD induced SARS-CoV-2 specific immune responses in BALB/c mice at one month post parenteral vaccination. **A.** Study design and vaccination timeline. **B.** Endpoint IgG subtype titers against SARS-CoV-2 r-RBD measured in serum collected from the vaccinated mice. n= 5. **C-D.** SARS-CoV-2 specific memory B cell (MBC) responses by ELISPOT analysis. **C.** Images of wells from MBC culture. Splenocytes were stimulated *in vitro* for 7 d with R848 plus rIL-2 and seeded onto ELISPOT plates coated with Ig capture Ab or SARS-CoV-2 RBD. Images of total IgG-antibody secreting cells (ASC), RBD-specific MBCs, and negative control (NC) wells are shown. **D.** Frequencies of SARS-CoV-2 RBD-specific ASCs per 10^6^ input cells in MBC cultures from the subject. n= 4. **E-F.** ELISPOT quantification of vaccine-specific T cells. Mouse splenocytes were *ex vivo* stimulated with overlapping peptide pools spanning SARS-CoV-2 S protein, α-CD3, or blank (negative control, NC) for 20 h. **E.** Images of wells from T cell culture. **F.** Spot forming cells (SFC) were measured by IFN-γ ELISPOT. Data are shown as # of SFC per 10^6^ splenocytes. n= 5. ** *P* < 0.01 or \**P* < 0.05 compared to mock group. ##*P* < 0.01 compared to alum-RBD group.

To assess the durability of mPSM-RBD-induced immunity, BALB/c mice were immunized i.d. with PBS (mock), mPSM-RBD (5 μg), or Alum-RBD (5 μg) on days 0 and 14. Longitudinal sera samples were collected over the course of 7 months to determine SARS-CoV-2-specific antibody responses (**Fig 3A**). mPSM-RBD vaccination triggered the production of SARS-CoV-2 RBD-binding IgG2a, IgG2b and IgG1 antibodies on day 10, which reached to the peak response around 4 weeks but remained high even at 7 months post vaccination. In contrast, RBD-binding IgG2a and IgG2b antibodies were barely detectable except for lower IgG1 responses in alum-RBD-vaccinated mice (**Fig 3B-D**). In addition, mPSM-RBD-vaccinated mice showed more than 100 times higher titers of RBD-binding total IgG 4.5 months post vaccination compared to mice treated with alum-RBD (**Fig S3A-B**). Furthermore, high Nab titers were detected at 1 month in the majority of mPSM-RBD-vaccinated mice and remained at a similar level 5 months later in all vaccinated mice; in comparison, NAb was barely detectable in any alum-RBD-vaccinated mice throughout the time (**Fig 3E**). Moreover, while both mPSM-RBD and alum-RBD vaccinations induced RBD-specific IgG^+^ B cell responses, there were 2.5-fold as many S -specific IgG^+^ splenic B cells and 1.5-fold as many SARS-COV-2-specific splenic Th1 cells in the mPSM-RBD group compared to the alum-RBD group 7 months post vaccination (**Fig 3F-J**). Both mPSM-RBD and alum-RBD-vaccinated mice showed higher SARS-COV-2 S-specific IgA^+^ splenic B cells than the mock group at the 7-month time point (**Fig S3C-D**). Taken together, parenteral vaccination with mPSM-RBD induced stronger and more durable SARS-CoV-2-specific IgG^+^ B cells, higher Nab titers, and Th1-prone immune responses than alum-RBD in mice.

**Figure 3.**
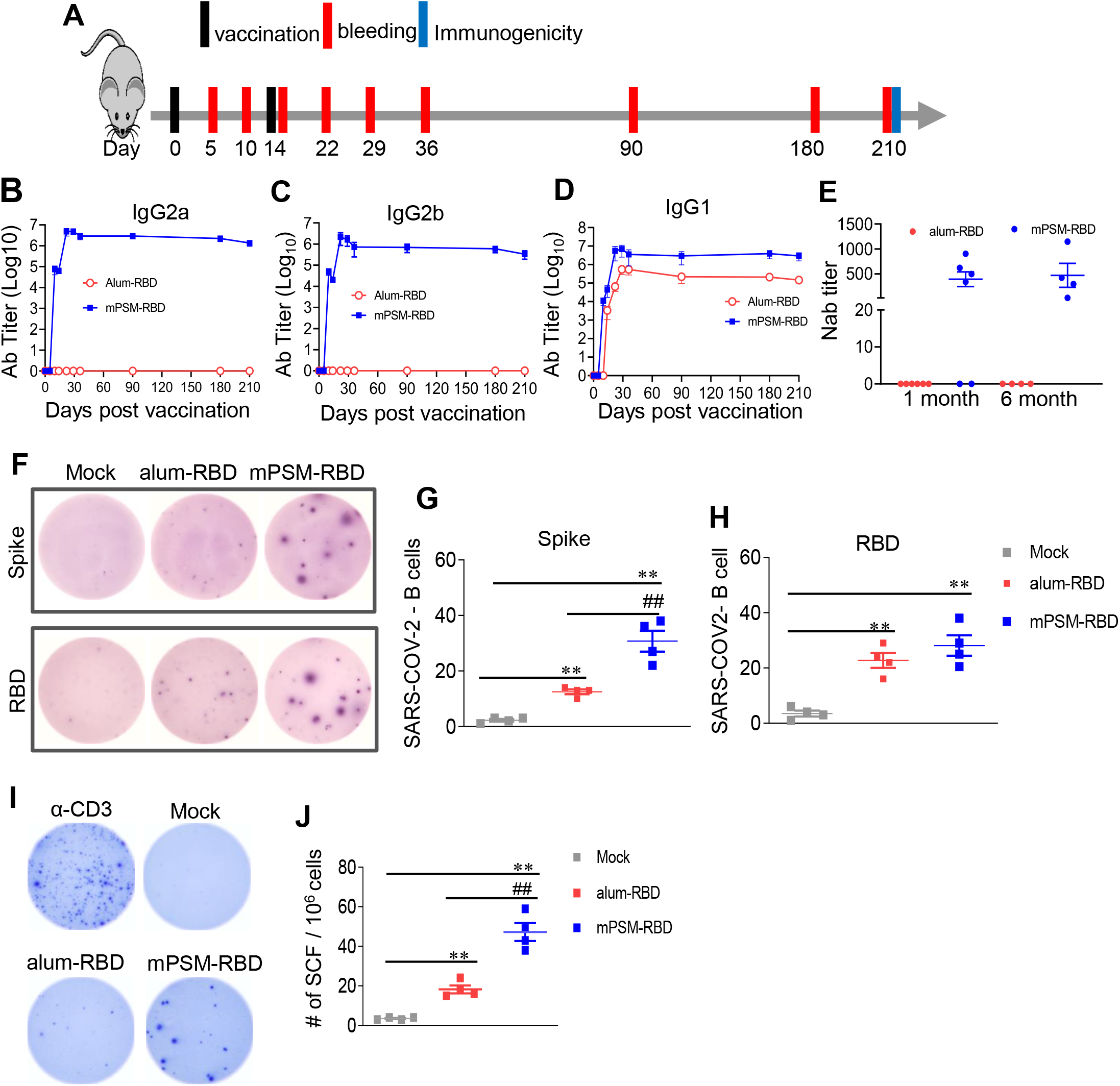
mPSM-RBD induced durable Type 1 prone protective immunity in mice. Six-week-old BALB/c mice were prime-boost immunized with mock (PBS), alum-RBD, or mPSM-RBD via i.d. route. **A**. Study design and vaccination timeline. **B-D.** Endpoint IgG subtype titers against SARS-CoV-2 r-RBD measured in serum collected at various time points after vaccination. n = 4. **E.** Serum SARS-CoV-2 neutralizing activity measured by plaque reduction neutralization test (PRNT). PRNT_80_ titers are shown, n = 4 or 6. **F-H.** SARS-CoV-2 specific memory B cell (MBC) responses by ELISPOT analysis at 7 months post vaccination. **F.** Images of wells from MBC culture. Frequencies of spike (**G**) or RBD (**H**) specific ASCs per 10^6^ input cells in MBC cultures from the subject. **I-J.** ELISPOT quantification of vaccine-specific splenic T cells at 7 months post vaccination. Mouse splenocytes were *ex vivo* stimulated with overlapping peptide pools spanning SARS-CoV-2 S protein, α-CD3, or blank for 20 h. **I.** Images of wells from T cell culture. **J.** Spot forming cells (SFC) were measured by IFN-γ ELISpot. Data are shown as # of SFC per 10^6^ splenocytes. n= 4. ** *P* < 0.01 compared to the mock group. ##*P* < 0.01 compared to alum-RBD group.

### mPSM-RBD provides more durable and potent protection against SARS-CoV-2 and Beta variant infection following single or two dose parenteral vaccination in mice

To assess the efficacy of mPSM-RBD in protecting the host against SARS-CoV-2 infection, BALB/c mice were vaccinated with alum-RBD (5 μg), mPSM-RBD (5 μg), or mock i.p. on day 0 and boosted with the same dose on day 21. At 1 month post vaccination, mice were i.n. challenged with 2 x 10^4^ PFU mouse-adapted SARS-CoV-2 strain CMA4 ^17^. Mice were euthanized two days after infection (**Fig S4A**). There were lower viral loads and attenuated levels of inflammatory cytokines, including CCL2, CCL7 and CXCL10 in the lung of mPSM-RBD group compared to the mock group. Alum-RBD-vaccinated mice also showed similar reductions on viral loads and inflammation in the lung (**Fig S4B-E**). In another study, mice were i.n. challenged with 2 x 10^4^ PFU of the mouse-adapted SARS-CoV-2 strain CMA4 at 4.5 months post vaccination. While mice in both mock and alum-RBD groups exhibited 10^2^ to 10^3^ PFU/ml viral loads in the lung tissues; no detectable viral titers were measured in the mPSM-RBD group at day 4 post infection (**Fig 4A-B**). In addition, lung inflammation was assessed by measurement of proinflammatory cytokines (IL-1β, IL-6) and chemokines (CCL2, CCL7, CXCL10) levels (**Fig 4C-H**). The mPSM-RBD-vaccinated mice had significantly reduced levels of inflammation compared to the mock and the alum-RBD group. Furthermore, to assess protective efficacy from a single dose vaccination, 6-8-week-old K18 hACE2 mice were treated i.p. with PBS (mock), alum-RBD (25 μg), or mPSM-RBD (25 μg). Mice were challenged i.n. with 4×10^3^ PFU of SARS-CoV-2 Beta variant 1 month post vaccination. While both alum-RBD and mPSM-RBD groups showed reduced viral loads in the lung compared to the mock group, mice in the mPSM-RBD group had 40% lower viral load in the lungs than those in the alum-RBD group (**Fig 4I, J**). In summary, these data showed that the mPSM-RBD vaccine triggered more durable and stronger protection against SARS-CoV-2 and Beta variant infection than the alum-RBD vaccine following single or two doses of parenteral vaccination.

**Figure 4.**
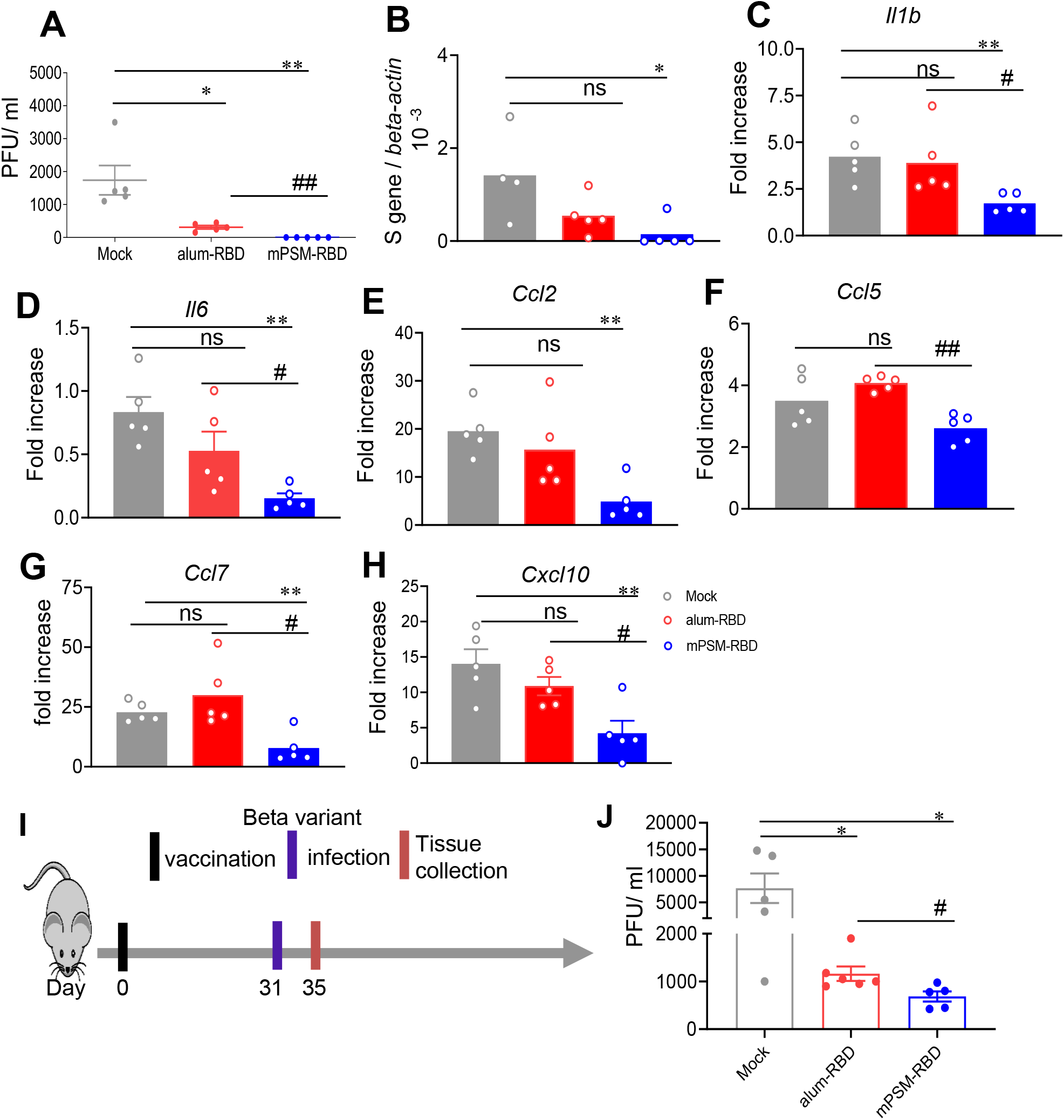
The protective efficacy of mPSM-RBD vaccine against SARS-CoV-2 and the Beta variant infection following single or two dose parenteral vaccination. **A-H.** Six-to eight-week-old BALB/c mice (n =5) were prime-boost immunized with mock (PBS), alum-RBD, or mPSM-RBD. At 4.5 months post vaccination, all mice were i.n. challenged with 2 x10^4^ PFU mouse-adapted SARS-CoV-2 CMA4. At day 4 post infection (pi), lung tissues were collected. (**A-B**) SARS-CoV-2 viral titers in lung tissues were measured by plaque (**A**) and Q-PCR (**B**) assays. **C-H**. Measurement of cytokine and chemokine levels in lung tissues by Q-PCR assays at day 4 post infection. Data are presented as the fold increase compared to naïve mice (means ± SEM). **I-J**. Six-week-old K18 ACE2 mice (n =5) were immunized once i.p. with mock (PBS), alum-RBD, or mPSM-RBD (25 ug). One month post vaccination, all mice were i.n. challenged with 4000 PFU SARS-CoV-2 Betha variant and lung tissues were collected at day 4 pi. **I.** Study design and timeline for vaccination and viral challenge. **J.** SARS-CoV-2 viral titers in lung tissues were measured by plaque assay. ** *P* < 0.01 or \**P* < 0.05 compared to mock group. #*P* < 0.05 compared to alum-RBD group.

### mPSM promotes nasal and airway epithelial cells uptake of SARS-CoV-2 RBD antigen; intranasal boost with mPSM-RBD triggers higher levels of SARS-CoV-2 -specific mucosal immune responses and protect the host against SARS-CoV-2 Delta variant infection

The magnitude of virus-specific T cells in the lung is known to be associated with better prophylaxis of COVID-19 patients ^18^. Mucosal vaccination is likely to be more effective in control of virus spread as it can enhance lung resident memory T cells compared to parenteral injection ^19^. To determine whether mPSM could also serve as an efficient carrier for mucosal delivery of SARS-CoV-2 antigen, we assessed RBD antigen uptake by the upper respiratory epithelial cells. Cy5-labeled mPSM-RBD was applied to treat human small airway epithelial cells (SAEC) and human nasal cell line RPMI2650, and intracellular particle trafficking was monitored. Microscopic analysis revealed that mPSM-RBD bound to both SAECs and RPMI2650 cells, with a higher binding affinity to SAECs based on the average number of particles in each cell type (Fig 5A). mPSM-RBD co-localized with early endosome (EEA1^+^, green) as soon as 0.5 h after incubation. After 2 h and 6 h incubation, mPSM-RBD vaccine was gradually released from the particles and reached the surrounding area inside the cells. These results suggest that mPSM can effectively deliver RBD antigen and promote its uptake by upper respiratory epithelial cells. Next, we assessed SARS-CoV-2-specific immune responses in BALB/c mice following primed i.p. with PBS (mock), RBD alone, m-PSM-RBD or alum-RBD (5 μg) on day 0 and boosted i.n. with the same dose on day 21 (Fig 5B). Blood, bronchoalveolar lavage fluids (BAL), lung and spleen tissues were collected on day 35. In the lung, there were stronger SARS-CoV-2-specific Th1 responses in mPSM-RBD group than the alum-RBD group, and both CD4^+^ and CD8^+^ T cells produced more IFNγ^-^ than the alum-RBD group (Fig 5C-E). While both alum-RBD and mPSM-RBD vaccinations triggered more RBD-specific IgA^+^ B cells in the lung compared to that of the mock group, the mPSM-RBD group produced at least 2-fold as many RBD-specific IgA^+^ B cells as those in the alum-RBD group (Fig 5F). In the spleen, the mPSM-RBD group showed elevated levels of IFNγ -production than the alum-RBD group. Among splenic T cells, CD8^+^ T cells, but not CD4^+^T cells, produced significantly more IFNγ in the mPSM-RBD group than the alum-RBD group (Fig S5A-B). SARS-CoV-2 specific IgA^+^ splenic B cells were also induced in the mPSM-RBD-vaccinated mice (Fig S5C). Furthermore, higher titers of RBD-binding IgA antibodies were detected in BAL and sera (Fig 5H, Fig S5D), as well as RBD-binding IgG1 and IgG2a antibodies in sera of mPSM-RBD-vaccinated mice compared to that of alum-RBD-vaccinated mice (Fig 5I, J). Lastly, to determine the effects of i.p/i.n. prime and boost with mPSM-RBD vaccine in host protection from SARS-CoV-2 variant infection, K18 hACE2 mice were vaccinated i.p. with PBS (mock), RBD (5 μg), m-PSM-RBD (5 μg), alum-RBD (5 μg) on day 0 and boosted i.n. with the same dose on day 21. Mice were then i.n. challenged with 1 x 10^4^ PFU of SARS-CoV-2 Delta variant at day 35. On day 4 post infection, plaque and Q-PCR assays showed that mPSM-RBD group had about 685-fold and 50-fold decrease in lung viral loads compared to the mock and alum-RBD groups, respectively (Fig 6A, B). In addition, the mPSM-RBD-vaccinated mice also showed significantly diminished levels of inflammatory cytokines in the lung compared to those in the mock group; in comparison, no difference was detected between the alum-RBD and mock groups (Fig 6C-E). In conclusion, these studies demonstrated that i.n. boost with mPSM-RBD vaccine triggers stronger lung resident B cell and Th1-type immune responses and IgA production and protects the host against SARS-CoV-2 Delta variant infection.

**Figure 5.**
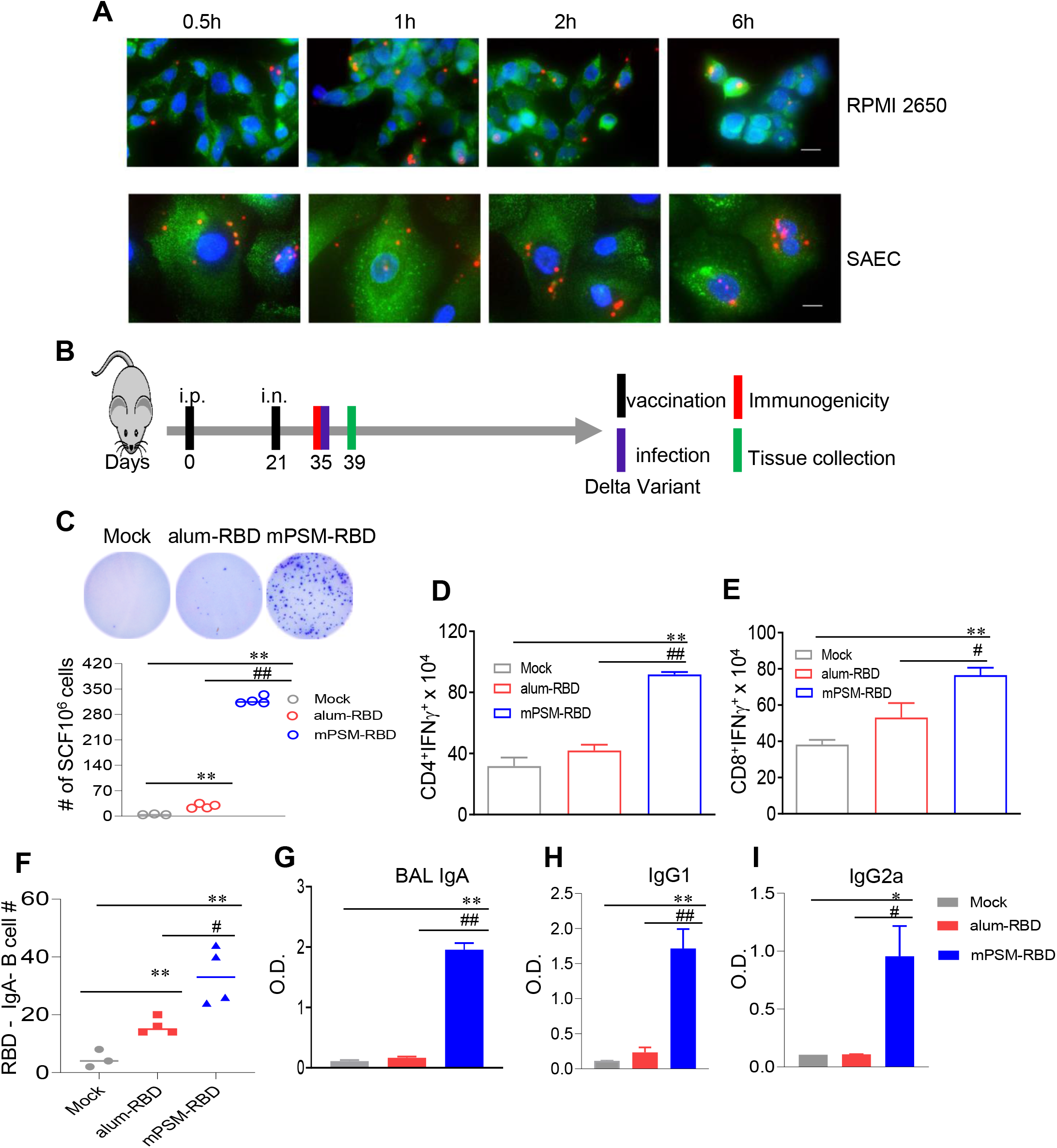
Parenteral and mucosal prime-boost vaccination promotes strong SARS-CoV-2 specific mucosal immune responses. (**A**) Fluorescence microscopic analysis on time-dependent uptake of vaccine particles in human small airway epithelial cells (SAE) and human nasal cell line RPMI2650. SAE cells and RPMI2650 cells were incubated with Cy5-labeled vaccine particles (red) for 0.5 h, 1 h, 2h and 6 h, respectively. Cells were then washed and stained with an anti-EEA1 antibody for early endosomes (green) and DAPI for nuclei (blue). Bar indicates 10 μm. (**B**) Study design and timeline for vaccination and viral challenge. Three groups of 6-8-week-old BALB/c or K18 ACE2 mice (n=5) were prime-boost immunized with mock (PBS), alum-RBD, or mPSM-RBD (5ug). At day 31 post vaccination, all mice were i.n. challenged with 1 x10^4^ PFU SARS-CoV-2 Delta variant. Four days after viral challenge, lung tissues were collected. **C-J.** Immunogenicity studies 1 month post vaccination in BALB/c mice. **C.** ELISPOT quantification of vaccine-specific lung T cells at 1 month post vaccination. Lung leukocytes were *ex vivo* stimulated with overlapping peptide pools spanning SARS-CoV-2 S protein, α-CD3, or blank for 20 h. Top panel. Images of wells from T cell culture. Lower panel. Spot forming cells (SFC) were measured by IFN-γ ELISPOT. Data are shown as # of SFC per 10^6^ cells. n= 3-4. **D-E.** Lung leukocytes were cultured *ex vivo* with S peptide pools for 5 h, and stained for IFN-γ, CD3, and CD4 or CD8. Total T cells were gated. Total number of IFN-γ^+^ CD4^+^ and CD8^+^ T cell subsets is shown. **F.** Lung leukocytes were stimulated *in vitro* for 7 days with R848 plus rIL-2 and seeded onto ELISPOT plates coated with SARS-CoV-2 RBD. Frequencies of SARS-CoV-2 RBD specific IgA secreting lung B cells per 10^6^ input cells in MBC cultures from the subject. n= 3-4. **G-I.** IgA titers in BAL (**G**) and IgG1, and IG2a subtypes in sera (**H-I**). ** *P* < 0.01 or \**P* < 0.05 compared to mock group. #*P* < 0.05 compared to alum-RBD group.

**Figure 6.**
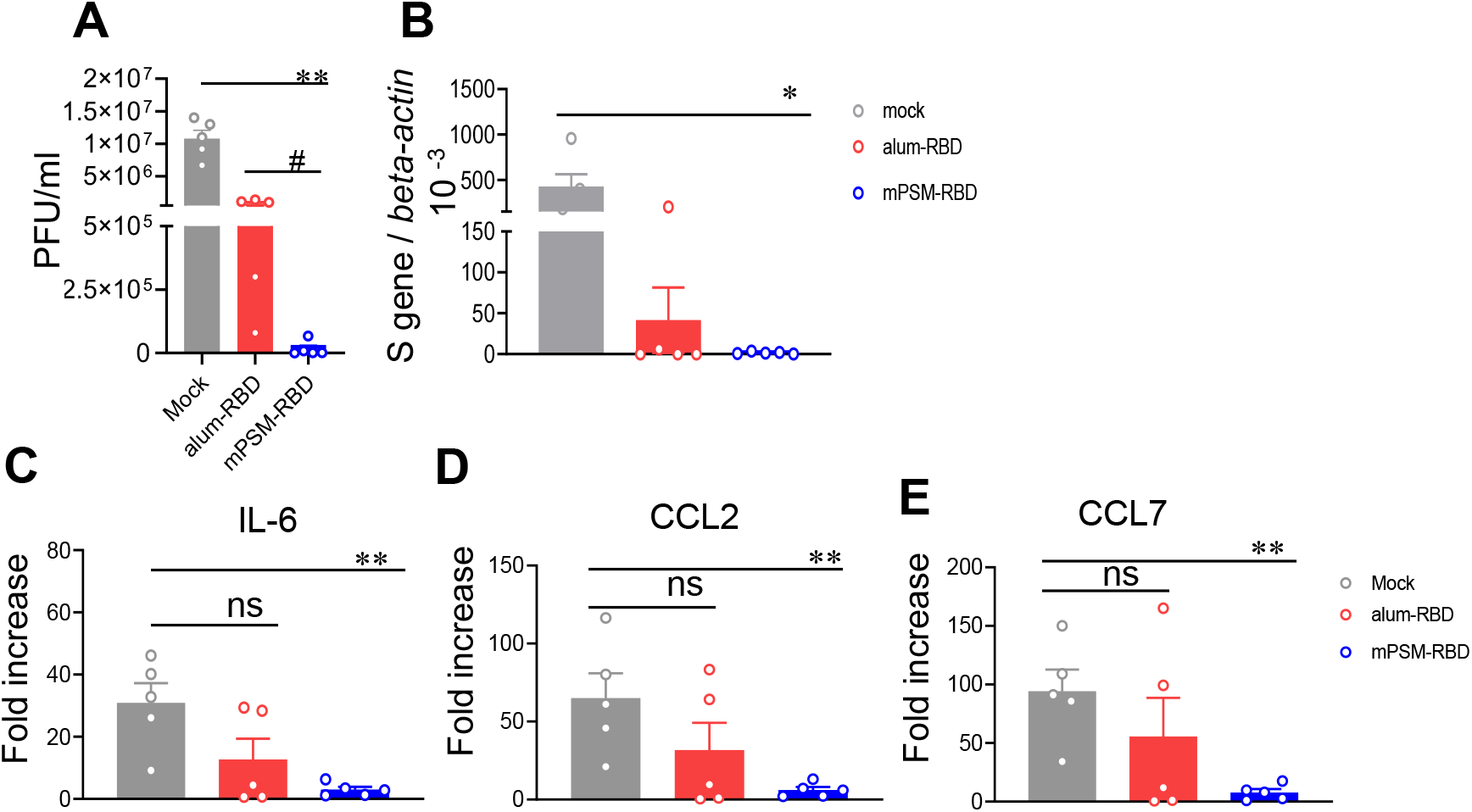
The protective efficacy of parenteral and mucosal prime-boost vaccination against SARS-CoV-2 Delta variant infection. As described in Fig. 5B, three groups of 6-8-week-old K18 ACE2 mice (n=5) were prime-boost immunized with mock (PBS), alum-RBD, or mPSM-RBD (5μg). At day 31 post vaccination, all mice were i.n. challenged with 1 x10^4^ PFU SARS-CoV-2 Delta variant. Four days after viral challenge, lung tissues were collected. (A) SARS-CoV-2 viral titers in lung tissues were measured by plaque (A) and Q-PCR (B) assays. C-G. Measurement of cytokine and chemokine levels in lung tissues by Q-PCR assays at day 4 post infection. Data are presented as the fold increase compared to naïve mice (means ± SEM). n= 5. ** *P* < 0.01 or \**P* < 0.05 compared to mock group. #*P* < 0.05 compared to alum-RBD group.

## DISCUSSION

B cell and antibody responses are critical for virus neutralization and disease control but are often of limited duration and breadth during SARS-CoV or SARS-CoV-2 infection ^20^. Variable and sometimes low NAb titers were reported in some convalescent COVID-19 patients, suggesting other immune factors contribute to the recovery from virus -induced diseases ^21^. T cells are known to play an important role in the clearance of SARS-CoV infection and host protection ^22-24^. Chen et al reported that SARS-CoV-2 infection caused a decrease in CD4^+^ and CD8^+^ T cell counts, and suppressed IFN-γ production by CD4^+^ T cells, which were associated with the disease severity of COVID-19 ^25^. Overall, balanced humoral and Th-1 directed cellular immune responses are important host protection against SARS-CoV-2 infection ^26^. The S protein, including RBD, can elicit highly potent and persistent NAbs and contain many T cell epitopes ^3^. Therefore, adjuvanted S or RBD protein subunit vaccines likely represent some of the most viable strategies for rapidly eliciting SARS-CoV-2 NAbs and CD4^+^ T cell responses of various qualities depending on the adjuvant used. Currently, the most commonly used adjuvants in human vaccination, such as alum, are effective at enhancing serum antibody titers, but not Th1 responses ^27,28^. A single dose vaccination with alum-formulated S protein induced a more Th2 prone response in mice ^29^. Optimized vaccine strategies include adding T helper epitope with RBD antigen or combing a TLR7/8 agonist with alum have been shown to effectively trigger strong humoral immunity supplemented with cellular immunity in mice and enhance NAb titers in various animal models ^30,31^. Here, we found that mPSM serves as a better adjuvant than alum for SARS-CoV-2 RBD subunit vaccines to elicit stronger and more durable Nabs, plus memory B cell and Th1 skewed immune responses in mice following parenteral and mucosal vaccination.

The PSMs contain 40-80 nm pores that can be loaded with nanoparticles, which were preferentially internalized by DCs over other types of phagocytic cells inside the body. Once inside the cells, PSM slowly degrades into non-toxic orthosilicic acid, a process that can last for as long as two weeks and the cargo inside the nanopores is gradually released ^32,33^. Thus, PSM acts as a reservoir for sustained release of antigen and other stimulatory factors, which offers the benefit of long-term stimulation of the APCs to trigger long-lasting immunity. Furthermore, PSM was previously reported to stimulate TRIF/MAVS-mediated pathways leading to activation of type I IFN responses ^14^. mPSMs, which includes PSM CpG and cGAMP elicits stronger innate cytokine response and more potent Th-1 biased immune responses, possibly due to the synergistic immune responses via multiple intracellular signaling pathways ^15^.

Intranasal immunization can lead to the induction of antigen-specific immunity in both the mucosal and systemic immune compartments ^19^. Delivery of antigens to the sites of infection and induction of mucosal immune responses in the respiratory tract, including IgA and resident memory B and T cells provides two additional layers of protection compared to systemic vaccination ^34^. Induction of mucosal IgA antibodies has been shown to help control several other respiratory viruses, such as SARS-CoV and RSV ^35-37^. Compared to IgG, IgA has been shown to more effectively control SARS-CoV-2 infection in the upper respiratory tract and nasal passages ^38^. Thus, mucosal vaccination appears to be more effective in control of SARS-CoV-2 infection and disease ^39,40^. Current delivery of the EUA SARS-CoV-2 vaccines is limited to parenteral injection, such as intramuscular route. In fact, less than 10% of the total 100 COVID-19 vaccines currently undergoing clinical trials utilizes the intranasal route ^34^. One of the challenges with intranasal subunit vaccines is that soluble antigens delivered to the nasal passages do not breach the epithelium but instead must be transported across the epithelial barrier by specialized microfold cells to present to DCs located underneath the epithelium^41^. Embedded in the submucosa is the nasal-associated lymphoid tissue (NALT), which is the first site for inhaled antigen recognition in the upper respiratory tract and includes B cells, T cells, and APCs. Formulation, size, and antigen type are important factors in mucosal vaccine development because they are critical for induction of mucosal immunity. Nanoparticles with size ranging from 20 to 200 nm ^42^ can serve as carriers for drug delivery to penetrate the mucosal surface and increase retention in the lung ^43^. mPSMs were previously reported to get trapped in endosomes for an extended amount of time, a process that benefits both DC activation and antigen processing ^14,15^. Here, we demonstrated that mPSM promotes the uptake of SARS-CoV-2 RBD antigens by nasal and airway epithelial cells. Moreover, due to relatively rapid turnover rates of mucosal antibody and lung-resident memory T cells, we applied a ‘prime and pull’ vaccination strategy ^44^. This begins with conventional parenteral vaccination to elicit systemic long-lived IgG response and broader repertoire memory B and T cells (prime), followed by an intranasal boost to recruit memory B and T cells to local lung resident memory cells and IgA production (pull) to mediate protective immunity. We found that the parenteral and mucosal prime-boost vaccination elicited robust SARS-CoV-2 -specific systemic and mucosal IgA and Th1-skewed immune responses, which protected mice from SARS-CoV-2 Delta variant infection.

Since the pandemic started, several major new variants have been identified as associated with increased viral transmission and disease severity in COVID-19 patients in the United Kingdom, South Africa, Brazil, United States, and more recently in India ^10,11^. Among them, the Beta variant, which was first identified in South Africa, has three mutations in the SARS-CoV-2 RBD protein, namely K417N, E484K and N501Y. The Delta variant carries seven mutations in S protein (T19R, G142D, del157/158, L452R, T478K, D614G, P681R) ^45^. Both Beta and Delta variants are of particular concern for their potential resistance to antibodies elicited by prior SARS-CoV-2 infection and/or vaccination ^46,47^. Furthermore, there is a potential concern of “vaccine-induced disease enhancement”, which was reported for certain SARS-CoV vaccine candidates ^12^ and inactivated RSV vaccines ^48^. The potential risk of ADE mediated by Fc-receptor could be increased due to waning immunity after vaccination and possibly mutations in the SARS-CoV-2 S protein ^49^. Due to the above concerns, the optimal COVID-19 vaccines will need to exhibit long-lasting immunity, be effective for various populations globally, and provide cross-reactive protection against emerging variants. Here, our results showed that the mPSM-RBD vaccine induced potent and durable Th-1 prone immune responses and protected mice from SARS-CoV-2, Beta and Delta variants infection. Furthermore, the mPSM-RBD vaccine did not cause toxicity in mice.

In conclusion, we have demonstrated that mPSM is a potent adjuvant for SARS-CoV-2 subunit vaccine and promotes intranasal delivery that triggers robust systemic and mucosal immunity. The m-PSM-based platform serves as a novel tool for the development of vaccines to effectively combat SARS-CoV-2 and other emerging RNA viruses or infectious pathogens that rely on Th1-mediated immunity.

## METHODS

### Vaccine preparation

To express and purify the RBD protein, the amino acid residues of 319 to 541 of SARS-CoV-2 S protein were cloned into the lentivirus vector, pCDH-CMV-MCS-EF1α-RFP (System Biosciences). To facilitate the secretion and purification of the protein, the first 19 residues of the S protein and a hexahistidine (6xHis) tag were fused at the N-terminal as a secretion signal and the C-terminal respectively. The vector was then packaged into lentivirus to transduce 293FT cells. RBD protein was purified from culture supernatant using His-Trap Excel nickel column (Cytiva). M-PSM was prepared to include 1 μg CpG ODN (Invivogen) 1826 and 0.5 μg cGAMP (Invivogen) in PSM (6 x10^7^ particles) as described previously ^14,15^. 25 ul of Imject Alum (ThermoFisher) was mixed with RBD protein 30 min before inoculation.

### Viruses

SARS-CoV-2 Beta variant, and Delta variant were obtained from the World Reference Center for Emerging Viruses and Arboviruses (WRCEVA) at the University of Texas Medical Branch (UTMB) and were amplified twice in Vero E6 cells. The generation of the mouse-adapted SARS-CoV-2 strain CMA4 was described in a recent study ^17^. The virus stocks for experiments were sequenced to ensure no undesired mutations in the S genes during the amplification in Vero E6 cells.

### Mice

6-week-old BALB/c mice, C57BL/(B)6 mice, and K18 hACE2 mice (stock #034860) were purchased from Jackson Lab. For vaccination, mice were inoculated intraperitoneally (i.p.), intradermally (i.d.), or intramuscularly (i.m.) with 5 to 25 μg RBD conjugated with mPSM or Alum on days 0, and 14 or 21. In some experiments, mice were i.p. primed on day 0 and boosted with the same dose on day 21 via i.n. inoculation. Vaccinated mice were challenged with 1 x 10^4^ PFU of SARS-CoV-2 CMA4, or Delta variant, or 4 x10^3^ PFU SARS-CoV-2 Beta variant. Infected mice were monitored twice daily for signs of morbidity. On days 2 or 4 post infection, mice were euthanized for tissue collection. All animal experiments were approved by the Animal Care and Use Committees at UTMB and Houston Methodist Academic Institute, respectively.

### *In vitro* DC maturation assay

Bone marrow (BM)-derived DCs were generated as described previously^14^. Briefly, BM cells isolated from BALB/c mice were cultured for 6 days in medium supplemented with granulocyte-macrophage colony-stimulating factor (GM-CSF) and IL-4 (Peprotech) to generate DCs. DCs were then treated with RBD alone or together with alum or mPSM at 37°C for 24 h. Cells were harvested and stained with antibodies for cell surface markers, including CD80 or CD86 antibodies (BioLegend), and acquired by a BD LSR II flow cytometer (BD Biosciences). Data were analyzed using FlowJo software (BD Biosciences).

### Antibody ELISA

Plates were coated with 1 μg/mL of purified SARS-CoV-2 RBD protein overnight at 4°C. Plates were blocked with 1% BSA for 45 min at 37°C. Diluted serum samples were added and incubated for 2 h at room temperature. This will be followed by a 1 h incubation with biotinylated HRP conjugated goat anti-mouse IgG subtype antibodies (Southern Biotech). 3,3’,5,5’ tetramethylbenzidine (TMB, BD Biosciences) were added to the well for 15 min and reactions were stopped by sulfuric acid. Absorbance at 450 nm and 570 nm were read and the absorbance at 570 nm was subtracted from the absorbance at 450 nm. Binding endpoint titers were determined using a cutoff value which is negative control+10x SD. In some experiments, ELISA plates were coated with 250 ng/well recombinant SARS-2 RBD protein (RayBiotech, USA) for overnight at 4°C. The plates were washed twice with phosphate-buffered saline, containing 0.05% Tween-20 (PBS-T) and then blocked with 8% FBS for 1.5 h. Sera or bronchoalveolar lavage (BAL) were diluted 1:40 to 1:100 or undiluted in blocking buffer and were added for 1 h at 37°C. Plates were washed five times with PBS-T. Goat anti-mouse IgG (Sigma, MO, USA), goat anti-mouse IgG1, Goat anti-mouse IgG2a, or goat anti-mouse IgG2b (Southern Biotech) coupled to alkaline phosphatase was added at a 1:1000 to 1:2000 dilutions for 1 h at 37°C. Color was developed with *p*-nitrophenyl phosphate (Sigma-Aldrich), and the intensity was read at an absorbance of 405 nm. For IgA measurement, goat anti-mouse IgA (Southern Biotech) coupled to horseradish peroxidase (HRP) was added as the secondary antibody at a 1:2000 dilution for 1 h at 37C, followed by adding TMB (3, 3, 5, 5’-tetramethylbenzidine) peroxidase substrate (Thermo Scientific) for about 15 min. The reactions were stopped by 1M sulfuric acid, and the intensity was read at an absorbance of 450 nm.

### Cytokine measurement by ELISA

TNF-α, IL-6, and IL-12p70 production were measured using the cytokine kits purchased from Invitrogen and following the instructions from the manufacturer.

### ACE2 inhibition assay

96-well plates were coated with 1 μg/mL of purified SARS-CoV-2 RBD protein overnight at 4 °C. Plates were washed with PBS with 0.05% TWEEN-20, followed by blocking with 1% BSA for 45 min at 37°C. Mouse sera were diluted at 1:100 in 1% BSA in PBS were incubated for 30 min at room temperature. Human recombinant ACE2-Fc-tag (Raybiotech) was then added at 1 μg/mL and incubated overnight at 4 °C, followed by incubation with 0.2 μg/mL anti-ACE2 (R&D) for 1 h at room temperature. Rabbit anti-goat IgG-HRP (Santa Cruz) at 1:8000 dilution was added for 30 min at room temperature. TMB was added for 15 min and the reaction was stopped by sulfuric acid. Absorbance at 450 nm and 570 nm were read and the absorbance at 570 nm was subtracted from the absorbance at 450 nm.

### Quantitative PCR (Q-PCR)

Viral-infected cells or tissues were resuspended in Trizol (Invitrogen) for RNA extraction. Complementary (c) DNA was synthesized by using a qScript cDNA synthesis kit (Bio-Rad). The sequences of the primer sets for cytokines, SARS-CoV-2 S gene and PCR reaction conditions were described previously ^50-52^. The PCR assay was performed in the CFX96 real-time PCR system (Bio-Rad). Gene expression was calculated using the formula 2^ ^-[C_t_(target gene)-C_t_(*β-actin*)]^ as described before ^53^.

### B cell ELISPOT assay

ELISPOT assays were performed as previously described ^54^ with some modifications. Briefly, splenocytes or lung leukocytes were stimulated with 1 μg/ml R848 and 10 ng/ml recombinant human IL-2 (Mabtech In, OH). Millipore ELISPOT plates (Millipore Ltd, Darmstadt, Germany) were coated with 100 μl SARS-CoV-2 RBD (RayBiotech, USA, 10 mg/ml) or rSARS-CoV-2 spike protein (R&D Systems). To detect total IgG or IgA expressing B cells, the wells were coated with 100 μL of anti-mouse IgG or IgA capture Ab (Mabtech In). Stimulated cells were harvested, and added in duplicates to assess total IgG, IgA ASCs, or SARS-CoV-2 specific B cells. The plates were incubated overnight at 37°C, followed by incubation with biotin-conjugated anti-mouse IgG (Mabtech In) for 2 h at room temperature, then 100 μL/well streptavidin-ALP was added for 1 h. Plates were developed with BCIP/NBT-Plus substrate until distinct spots emerge, washed with tap water, and scanned using an ImmunoSpot 6.0 analyzer and analyzed by ImmunoSpot software (Cellular Technology Ltd).

### IFN-γ ELISPOT

Millipore ELISPOT plates (Millipore Ltd) were coated with anti-IFN-γ capture Ab (Cellular Technology Ltd) at 4°C overnight. Splenocytes or lung leukocytes were stimulated in duplicates with SARS-CoV-2 S peptide pools (2 μg/ml, Miltenyi Biotec) for 24 h at 37°C. Cells were stimulated with anti-CD3 (1 μg/ml, e-Biosciences) or medium alone were used as controls. This was followed by incubation with biotin-conjugated anti-IFN-γ (Cellular Technology Ltd) for 2 h at room temperature, and then alkaline phosphatase-conjugated streptavidin for 30 min. The plates were washed and scanned using an ImmunoSpot 6.0 analyzer and analyzed by ImmunoSpot software to determine the spot-forming cells (SFC) per 10^6^ splenocytes.

### Intracellular cytokine staining (ICS)

Splenocytes or lung leukocytes were incubated with SARS-CoV-2 S peptide pools (1μg/ml, Miltenyi Biotec) for 24 h. BD GolgiPlug (BD Bioscience) was added to block protein transport at the final 6 h of incubation. Cells were stained with antibodies for CD3, CD4, or CD8, fixed in 2% paraformaldehyde, and permeabilized with 0.5% saponin before adding anti-IFN-γ, or control rat IgG1 (e-Biosciences). Samples were processed with a C6 Flow Cytometer instrument. Dead cells were excluded based on forward and side light scatter. Data were analyzed with a CFlow Plus Flow Cytometer (BD Biosciences).

### Immunofluorescence staining

SAEC and RPMI2650 cells were seeded in 8-well chamber slides at a density of 3 x 10^4^ cells per well and cultured overnight. Fluorescent vaccine particles were prepared using Cy5 labeled CpG ODN, and then incubated with cells at the ratio of 10 to 1 between mPSM to cells for 6 h. After incubation, cells were washed with PBS twice, fixed with 4% paraformaldehyde at room temperature for 15 min, and permeabilized with 0.1% tween-20 for 15 min. After blocking with 1% BSA plus 5% FBS, cells were incubated with anti-EEA1 antibody (1:500, Abcam) at 4°C overnight, followed by staining with AF488 -labeled goat anti-rabbit secondary antibody (1:1000 dilution, ThermoFisher) at room temperature for 2 h. Finally, nuclei were stained with 0.5 μg/mL DAPI for 15 min.

### mNG SARS-CoV-2 reporter neutralization assay

The mNG SARS-CoV-2 reporter neutralization assay was performed using a previous method ^55^ with some modifications. Vero CCL-81 cells (1.2 ×10^4^) in 50 μl of DMEM containing 2% FBS were seeded in each well of black μCLEAR flat-bottom 96-well plate (Greiner Bio-one™). The cells were incubated overnight at 37°C with 5% CO_2_. On the next day, each serum in duplicate was two-fold serially diluted in DMEM with 2% FBS and incubated with mNG SARS-CoV-2 at 37°C for 1 h. The virus-serum mixture was transferred to the Vero CCL-81 cell plate with the final multiplicity of infection (MOI) of 0.5. For each serum, the starting dilution was 1/50 with nine two-fold dilutions to the final dilution of 1/ 12800. After incubating the infected cells at 37°C for 16-24 h, 25 μl of Hoechst 33342 Solution (400-fold diluted in Hank’s Balanced Salt Solution; Gibco) was added to each well to stain the cell nucleus. The plate was sealed with Breath-Easy sealing membrane (Diversified Biotech), incubated at 37°C for 20 min, and quantified for mNG fluorescence on CytationTM 7 (BioTek). The raw images (1 picture per well) were acquired using 4 × objective. Infection rates were determined by dividing the mNG positive cell number by total cell number (indicated by nucleus staining). Relative infection rates were obtained by normalizing the infection rates of serum-treated groups to those of non-serum-treated controls. The curves of the relative infection rates versus the serum dilutions (Log_10_ values) were plotted using Prism 8 (GraphPad). A nonlinear regression method was used to determine the dilution fold that neutralized 50% of mNG fluorescence (NT_50_).

### Plaque assay

Vero E6 cells were seeded on 6-well plates and incubated at 37 °C, 5% CO_2_ for 16 h. Lung tissue homogenates in 0.2 ml volumes were used to infect the cells for 1 h. After the incubation, the overlay medium containing MEM with 2% FBS, 1% penicillin-streptomycin, and 1.6% agarose was added to the infected cells. Plates were stained with neutral red (Sigma-Aldrich) and plaques were counted to calculate viral titers expressed as PFU/ml.

### Statistical analysis

Values for viral load, cytokine production, antibody titers, and T cell response experiments were compared using Prism software (GraphPad) statistical analysis and were presented as means ± SEM. *P* values of these experiments were calculated with a non-paired Student’s t test.

### Supplementary Methods

#### Serum biochemistry assay

Serum samples were tested for alanine aminotransferase (ALT), albumin (ALB), alkaline phosphatase (ALP), amylase (AMY), calcium (CA), creatinine (CRE), globulin (GLOB), glucose (GLU), phosphorus (PHOS), potassium (K+), Sodium (NA+), total bilirubin (TBIL), total protein (TP), and urea nitrogen (BUN) Biochemistry Panel Plus analyzer discs (Abaxis).

## ACKNOWLEDGEMENTS

This work was supported in part by NIH grants R01AI127744 (T.W.), R01 AI116812 (to X.B), R21AG069226 (X.B), U54CA210181 (H.S.), a Fast Grant from Emergent Ventures at the Mercatus Center (T.W.), and a Pilot Grant from the Institute for Human Infections &Immunity (IHII) at UTMB (X.B.). We thank Gang Li for technical assistance on the cloning of RBD protein and Dr. Linsey Yeager for assisting in manuscript preparation.

## AUTHOR CONTRIBUTIONS

A.A., Q.S., B.W., J.Z., J.M., S.R.O., and W.W. performed the experiments. X.B., P.Y.S., H.S., and T.W., designed the experiment. X.X., P.Y.S., and P.V.A. provided critical reagents, A.A., Q.S., J.Z., J.M., and T.W. analyzed the data. T.W. wrote the initial draft of the manuscript and other coauthors provided editorial comments.

## SUPPLEMENTARY FIGURE LEGENDS

**Supplementary Figure 1. mPSM serves as a potent but safe adjuvant for SARS-CoV-2 RBD vaccine. A**. Activation of cell surface CD80 expression in BMDCs 24 h after treatment with mPSM-RBD, Alum-RBD, RBD or mock. CD80 expression was measured by flow cytometry analysis. One representative image was shown. **B-H.** The pathogenic effects of mPSM-RBD in mice. Six-to eight-week-old female BALB/c mice (n =5) were i.p. inoculated with mPSM-RBD (5 μg) or PBS (mock). Sera were collected at 24 h post-vaccination for analysis using Biochemistry Panel Plus analyzer discs (**B-E**, Abaxis) or proinflammatory cytokine levels by Q-PCR (**F-H**). Data are presented as the fold increase compared to naïve mice (means ± SEM). n= 5.

**Supplementary Figure 2. mPSM-RBD induces SARS-CoV-2 specific immune responses in C57BL/6 mice one month post parenteral vaccination. A-B**. Endpoint IgG subtype titers against SARS-CoV-2 rRBD measured in serum collected from vaccinated mice. n= 5. **C-D.** SARS-CoV-2 specific memory B cell (MBC) responses by ELISPOT analysis. **C.** Images of wells from MBC culture. Splenocytes were stimulated *in vitro* for 7 d with R848 plus rIL-2 and seeded onto ELISPOT plates coated with Ig capture Ab or SARS-CoV-2 RBD. Images of total ASCs, RBD specific MBCs, and negative control (NC) wells are shown. **D.** Frequencies of SARS-CoV-2 RBD specific ASCs per 10^6^ input cells in MBC cultures from the subject. n= 4. **E-F.** ELISPOT quantification of vaccine-specific T cells. Mouse splenocytes were *ex vivo* stimulated with overlapping peptide pools spanning SARS-CoV-2 S protein, α-CD3, or blank (NC) for 20 hours. **E.** Images of wells from T cell culture. **F.** Spot forming cells (SFC) were measured by IFN-γ ELISPOT. Data are shown as # of SFC per 10^6^ splenocytes. n= 4. ** *P* < 0.01 compared to mock group. ##*P* < 0.01 compared to alum-RBD group.

**Supplementary Figure 3. mPSM-RBD induces durable Type 1 prone immune responses following parenteral vaccination. A-B**. IgG responses 4.5 months after vaccination. Six-week-old BALB/c mice were prime-boost immunized with mock (PBS), alum-RBD, or mPSM-RBD via i.p. route. **A**. O.D. values by ELISA. **C-D.** SARS-CoV-2 specific IgA expressing memory B cell (MBC) responses by ELISPOT analysis at 7 months post vaccination. **C.** Images of wells from MBC culture. Frequencies of RBD (**D**) specific ASCs per 10^6^ input cells in MBC cultures from the subject. ** *P* < 0.01 or \**P* < 0.05 compared to mock group. ##*P* < 0.01 compared to alum-RBD group.

**Supplementary Figure 4. The protective efficacy of mPSM-RBD vaccine against SARS-CoV-2 infection one month after parenteral vaccination.** Six-to eight-week-old BALB/c mice (n =5) were prime-boost immunized with mock (PBS), alum-RBD, or mPSM-RBD. One month post vaccination, all mice were i.n. challenged with 2 x10^4^ PFU mouse-adapted SARS-CoV-2 CMA4. At day 2 post infection (pi), lung tissues were collected. **A**. Study design and vaccination timeline. **B.** SARS-CoV-2 viral titers in lung tissues were measured by Q-PCR assay. **C-E**. Measurement of chemokine levels in lung tissues by Q-PCR assays at day 2 post infection. Data are presented as the fold increase compared to naïve mice (means ± SEM). ** *P* < 0.01 or \**P* < 0.05 compared to mock group.

**Supplementary Figure 5. Parenteral and mucosal prime-boost vaccination induced strong SARS-CoV-2 specific systemic immune responses.** Three groups of 6-8-week-old BALB/c were i.p. and i.n. prime-boost immunized with mock (PBS), alum-RBD, or mPSM-RBD (5ug). At day 31 post vaccination, blood and spleen tissues were collected for immunogenicity studies. A. ELISPOT quantification of vaccine-specific splenic T cells at 1 month post vaccination. Splenocytes were *ex vivo* stimulated with overlapping peptide pools spanning SARS-CoV-2 S protein, α-CD3, or blank for 20 h. Top panel. Images of wells from T cell culture. Lower panel. Spot forming cells (SFC) were measured by IFN-γ ELISPOT. Data are shown as # of SFC per 10^6^ cells. n= 3-4. B. Splenocytes were cultured *ex vivo* with S peptide pools for 5 h, and stained for IFN-γ, CD3, and CD4 or CD8. Total T cells were gated. Total number of IFN-γ^+^ CD4^+^ and CD8^+^ T cell subsets is shown. C. Splenocytes were stimulated *in vitro* for 7 d with R848 plus rIL-2 and seeded onto ELISPOT plates coated with SARS-CoV-2 RBD. Frequencies of SARS-CoV-2 RBD specific IgA secreting splenic B cells per 10^6^ input cells in MBC cultures from the subject. n= 3-4. D. IgA response in sera (I-J). ** *P* < 0.01 or \**P* < 0.05 compared to mock group. ##*P* < 0.01 or #*P* < 0.05 compared to alum-RBD group.

