## Supplementary figures and images for "A modified porous silicon microparticle promotes mucosal delivery of SARS-CoV-2 antigen and induction of potent and durable systemic and mucosal T helper 1 skewed protective immunity"

### Supplementary figures

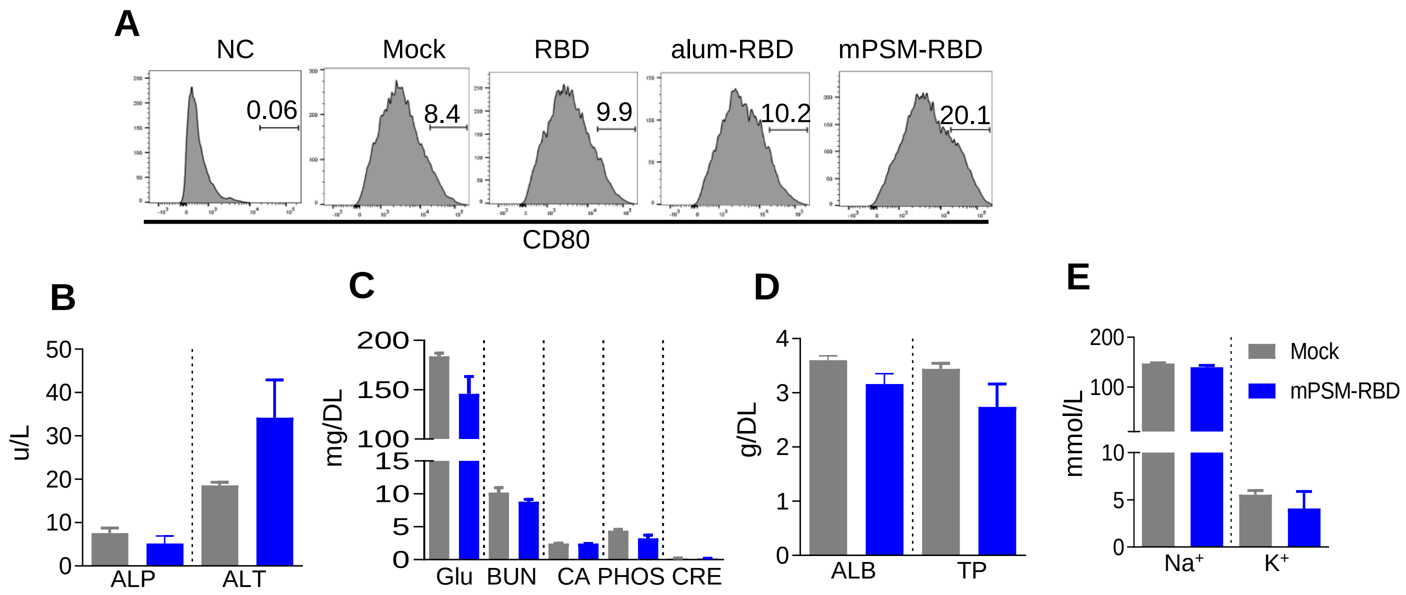

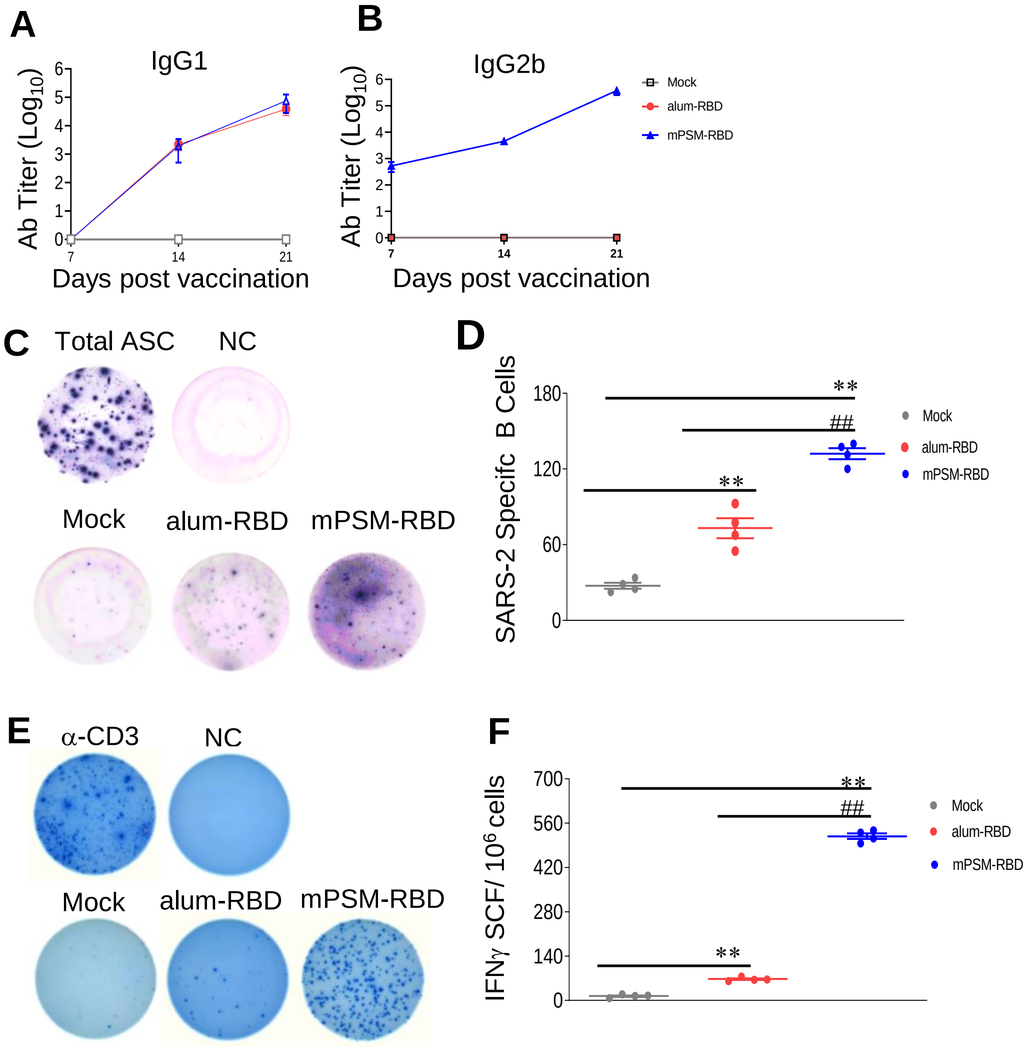

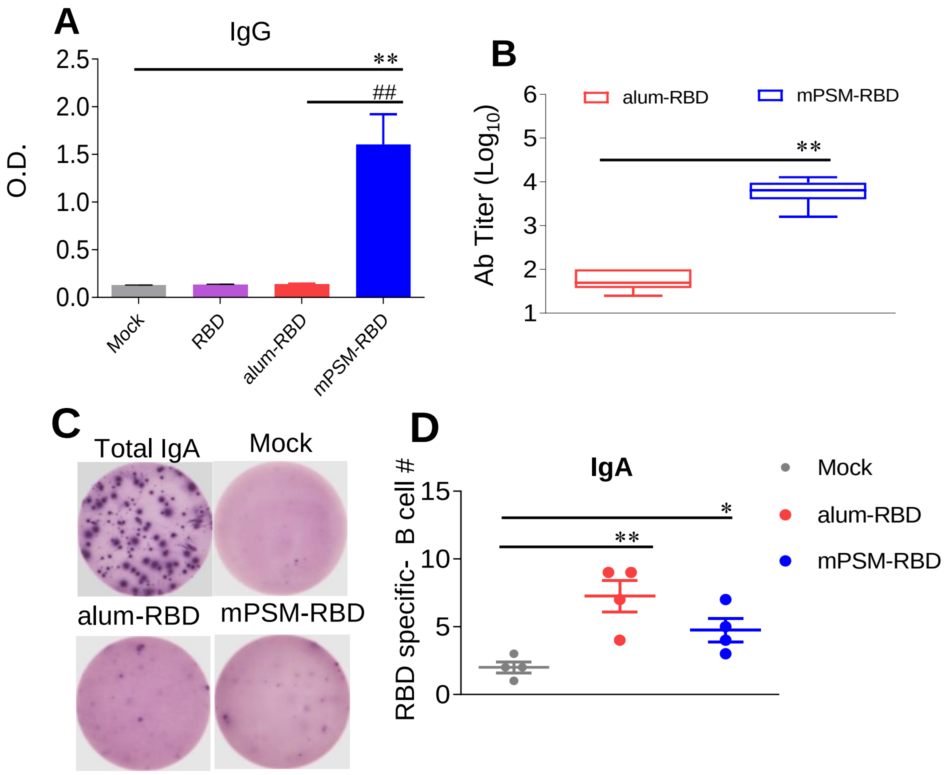

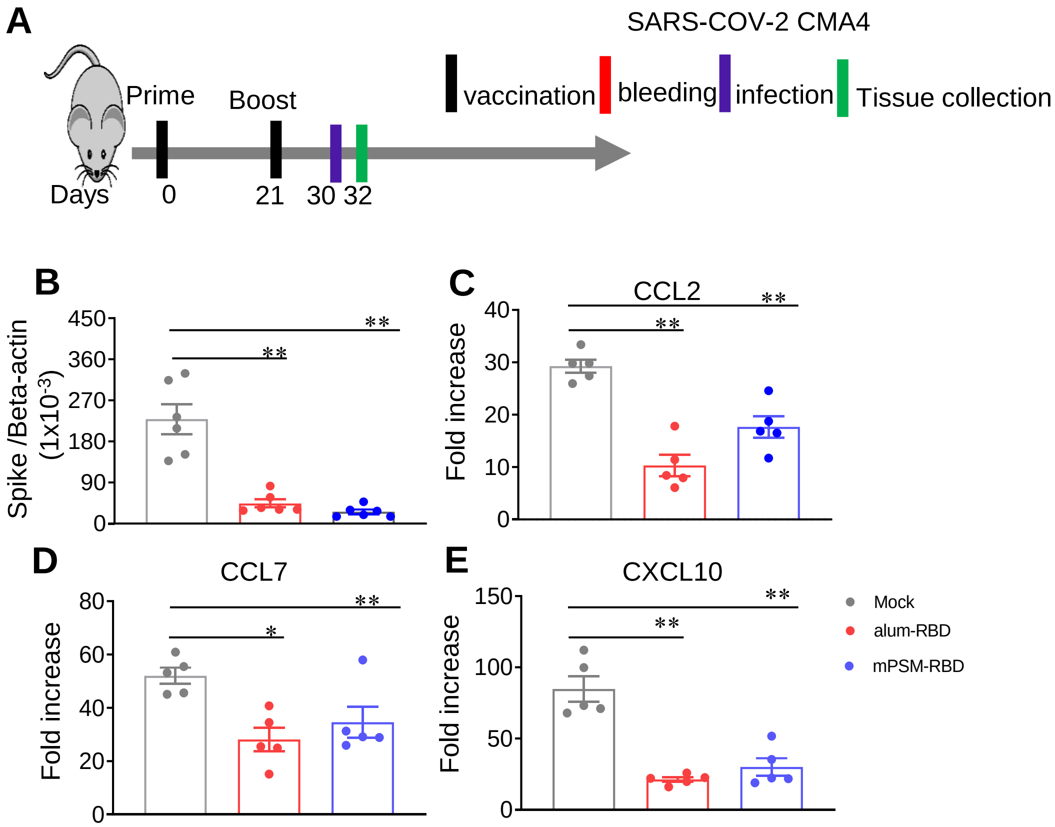

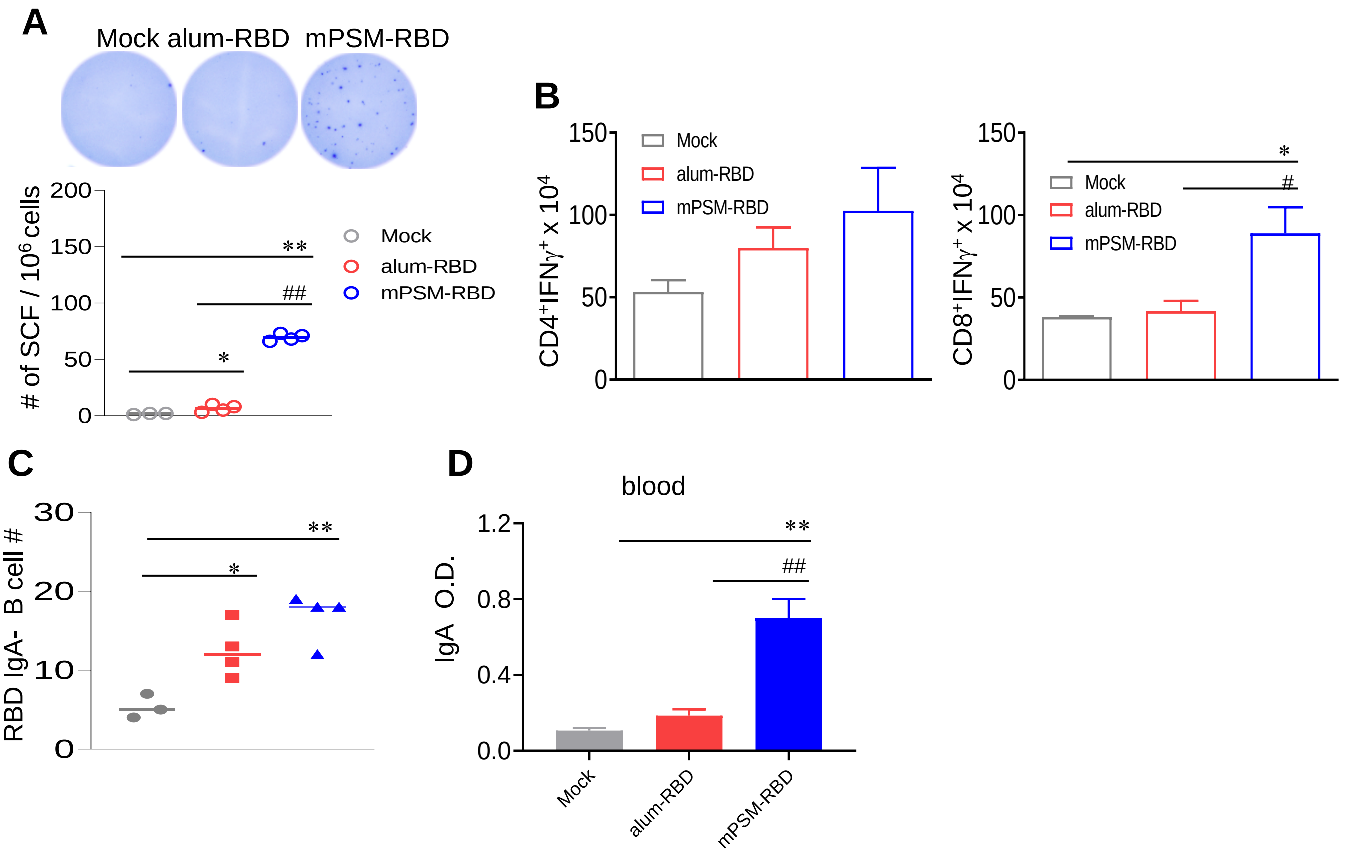
